# *In vivo* monitoring of plant small GTPase activation using a Förster resonance energy transfer biosensor

**DOI:** 10.1101/249938

**Authors:** Hann Ling Wong, Akira Akamatsu, Qiong Wang, Masayuki Higuchi, Tomonori Matsuda, Jun Okuda, Ken-ichi Kosami, Noriko Inada, Tsutomu Kawasaki, Shingo Nagawa, Li Tan, Yoji Kawano, Ko Shimamoto

## Abstract

Small GTPases act as molecular switches that regulate various plant responses such as disease resistance, pollen tube growth, root hair development, cell wall patterning and hormone responses. Thus, to monitor their activation status within plant cells is believed to be the key step in understanding their roles. We have established a plant version of a Förster resonance energy transfer (FRET) probe called Ras and interacting protein chimeric unit (Raichu) that can successfully monitor activation of the rice small GTPase OsRac1 during various defence responses in rice cells. Here, we describe the protocol for visualizing spatiotemporal activity of plant Rac/ROP GTPase in living plant cells, transfection of rice protoplasts with *Raichu-OsRac1* and acquisition of FRET images. Our protocol should be widely adaptable for monitoring activation for other plant small GTPases and for other FRET sensors in various plant cells.

## INTRODUCION

### Background

Scientists have wanted for many years to observe events in living cells at the molecular level, and fluorescent proteins offer powerful tools for doing so. Förster Resonance Energy Transfer (FRET) is a physical phenomenon that can be sensitive to changes in molecular conformation and association typically in the 1-10 nm range (*1*, *2*). In FRET applications, a pair of donor and acceptor fluorescent proteins is used to monitor a variety of biological events including protein-protein interactions, protein kinase activities, post-translational modifications and the dynamics of second messengers (*3–6*). When the emission spectrum of a donor fluorescent protein overlaps with the absorption spectrum of an acceptor fluorescent protein, and the distance between the two proteins is less than 10 nm, non-radiative transfer of energy occurs from the donor to the acceptor. Currently, cyan-emitting fluorescent protein (CFP) and yellow-emitting fluorescent protein (YFP) are widely used as the donor and acceptor pair for FRET analyses.

Small GTPases exhibit both GDP/GTP-binding activity and GTPase activity, and work as molecular switches by cycling between GDP-bound inactive and GTP-bound active forms. The Ras superfamily GTPases contain five highly conserved G-boxes (G1-G5) (*7*, *8*). G1, G3, G4 and G5 play vital roles in binding to GTP/GDP and in GTPase activity, while the G2 box is an effector domain that is important for binding to downstream effector proteins. The C-terminal polybasic region and post-translational modification sites contribute to subcellular localization and function.

The Ras superfamily is classified structurally and functionally into at least five families: Ras, Rho, Rab, Sar1/Arf and Ran (*8*). The Rho family in animals is further split into three subfamilies, namely Rho, Rac and Cdc42. In plants, the Rho family converges into a single subfamily that is distinct from the Rho, Rac and Cdc42 subfamilies in animals (*9*), although, functionally, the plant Rho subfamily is most closely related to the animal Rac subfamily. Thus, these plant Rho subfamily GTPases are called Rac or ROP (Rho of plant or Rho-related GTPases from plants) proteins (*10*, *11*). The ratio of active GTP-bound to inactive GDP-bound forms of a Rac/ROP GTPase is controlled by the activity of three factors (*12*): GTPase-activating proteins (GAPs) act as negative regulators by accelerating the intrinsic GTPase activities of Rac/ROP and converting it to an inactive GDP-bound Rac/ROP; guanine nucleotide dissociation inhibitors (GDIs) suppress the exchange of GDP for GTP; guanine nucleotide exchange factors (GEFs) promote the release of GDP from Rac/ROP and, since the intracellular concentration of GTP is much higher than that of GDP, thereby facilitate the binding of GTP to Rac/ROP. GTP-bound Rac/ROP associates with downstream effector proteins to trigger a variety of cellular responses.

### Comparison with other methods

Rac/ROP family small GTPases are master regulators, controlling various signalling systems in plants such as those governing pollen tube growth, root hair development, auxin signaling, and disease resistance (*13–16*). Therefore, monitoring the activation state of a Rac/ROP GTPase within living cells is critical to further understanding its functions. Until the 1990s, activation states of small GTPases were measured *in vivo* by labelling cells with inorganic [^32^P] phosphate, followed by immunoprecipitation of small GTPases and thin-layer chromatography to obtain quantitative data for their associated GDP and GTP levels. However, this method required not only large amounts of radioisotope but also a time-consuming procedure. Currently, two alternative non-radioactive techniques are available for monitoring the *in vivo* activation of Rac/ROP: a PAK CRIB pull-down assay, and FRET sensors (*5*, *17–19*). These methods exploit the selective interaction between the Cdc42/Rac interactive binding (CRIB) domain of the Rac-effector PAK1 (PAK CRIB) and plant and animal Rac GTPases. PAK CRIB shows a high affinity only for the active GTP-bound form of Rac, and not for the inactive GDP-bound form (*17*, *19*). PAK CRIB binding suppresses the intrinsic GTPase activity of Rac, and this function provides a useful tool for isolating active GTP-bound forms of Rac/ROP from crude cell lysates and for monitoring the activation state of Rac/ROP in living cells. Currently, kits using GST-tagged PAK CRIB for a PAK CRIB pull-down assay are available from several manufacturers (e.g., Cytoskeleton). The subsequent immunoblotting assay can quantify Rac/ROP activation in cells by determining the amount of activated Rac/ROP. This is a widely used method for the semi-quantitative measurement of Rac/ROP activity from plants to animals (*17*, *18*, *20*). However, the PAK CRIB pull-down assay only provides a snapshot, and cannot detect the spatial and temporal dynamics of intracellular signaling in living cells. To solve this problem, FRET biosensors for small GTPases have been developed.

### Development of the protocol

FRET biosensors that have been developed to visualize small GTPase activity in living cells can be classified into two types: intramolecular (or unimolecular) and intermolecular (or bimolecular) biosensors. Both types exploit the ability of two different proteins to interact with each other when one of them is activated. Intramolecular biosensors have both donor and acceptor fluorescent proteins in one molecule, whereas a donor and an acceptor fluorescent protein are expressed separately in intermolecular biosensors. The first report on *in vivo* monitoring of a small GTPase involved intermolecular biosensors, namely the combination of animal cultured cells expressing green fluorescent protein (GFP)-Rac as the donor and microinjection of PAK CRIB labeled with Alexa 546 dye as the acceptor, which binds selectively to the GTP-active form of Rac (*21*). The authors were able to show clearly the spatial control of growth factor-induced Rac activation in a membrane ruffling area, as well as a gradient of activation, at the leading edge of motile cells. In plants, an intermolecular FRET biosensor was built to analyze ROP2 activity. With this sensor, which used a downstream effector protein of the small GTPase ROP2, CFP-RIC4 (donor), and YFP-ROP2 (acceptor), it was shown that ROP2 is preferentially activated in lobe-forming regions of the cell cortex (*22*).

However, there are several disadvantages in systems involving intermolecular biosensors. For example, to obtain reliable data from FRET analyses, we have to control the expression levels of donors and acceptors (*23*). The expression level of the acceptor should be higher than that of the donor; moreover, a high proportion of activated donors must associate with acceptors to obtain sufficiently high signal/noise ratios to prevent FRET signals from being buried under noise. In plants, particle bombardment of plasmids into leaf epidermal pavement cells of *Arabidopsis* is often carried out for FRET analyses, but it is technically difficult to obtain appropriate levels of expression of donors and acceptors in *Arabidopsis* using intermolecular biosensors. Moreover, excessive expression of acceptors can cause abnormal activation or inhibition of downstream molecules.

To overcome these disadvantages, Matsuda and his colleagues developed excellent intramolecular FRET biosensors for small GTPases in animal cells, collectively naming them Raichu (Ras superfamily and interacting protein chimeric unit). Raichu was initially developed to study activation of the small GTPases Ras and Rap1 following growth factor stimulation in animal cells (*5*, *24*). The original Raichu contains a donor (cyan-emitting fluorescent protein; CFP), an acceptor (yellow-emitting fluorescent protein; YFP), and the Ras-binding domain of Raf (RBD), which is a downstream effector and binds specifically to active Ras. Therefore, the molar ratio of the individual component proteins is the same irrespective of expression level (Fig. 1a). Accordingly, this intramolecular FRET biosensor eliminates the problem of variability in the expression levels of donor and acceptor fluorescent proteins and is an ideal sensor for monitoring the activation states of small GTPases.

**Figure 1.**
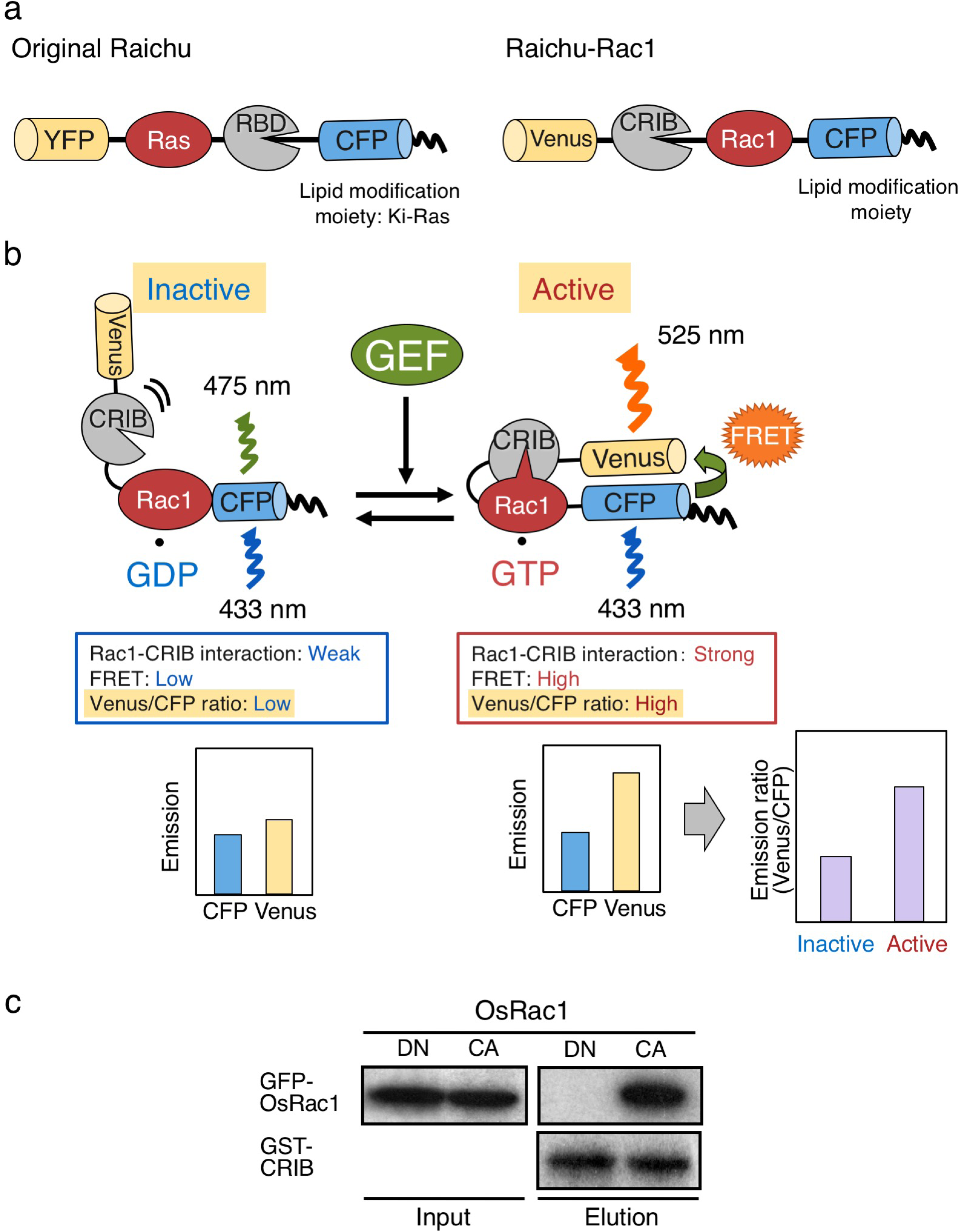
Mechanism of Raichu-OsRac1 FRET sensor. **a,** Schematic representation of original Raichu and Raichu-Rac1 intramolecular FRET biosensors. **b,** Raichu-Rac1 consists of the fluorescent protein Venus (yellow), the CRIB domain of PAK (grey), the small GTPase Rac1 (red) and the fluorescent protein CFP (cyan). When OsRac1 is bound to GDP, the intramolecular association between the CRIB domain of PAK is weak, and fluorescence of 475 nm thus emanates from CFP upon excitation at 433 nm. When OsRac1 is bound to GTP, intramolecular interaction between the PAK CRIB domain and OsRac1 brings CFP and Venus into close proximity, causing FRET and fluorescence of Venus at 525 nm. **c,** Binding specificity of the PAK CRIB domain for OsRac1 in an *in vitro* binding assay. Sepharose-immobilized GST-CRIB was incubated with extracts from HEK293 cells expressing GFP-CA-OsRac1 or GFP-DN-OsRac1. After washing, bound proteins were eluted by the addition of SDS-PAGE sample buffer. The eluted fractions were immunoblotted with anti-GFP (Roche) (upper panels) and anti-GST (Santa Cruz) (lower panel) antibodies.

Subsequently, Raichu and its variants have become well-established tools for visualizing the activation of various small GTPases, including Rac1, Cdc42, RhoA, Ral, TC10 and Rab5 in animals (*15*, *25*). Raichu-Rac1, one of the variants of Raichu, is composed of the yellow-emitting fluorescent protein Venus, the small GTPase human Rac1, a linker, the CRIB domain of PAK, CFP, and the C-terminal polybasic region and post-translational modification site of KiRas at the C terminus (Fig. 1 a). In the GDP-bound inactive form of Raichu-Rac1, PAK CRIB does not bind to Rac1 and the donor CFP remains remote from the acceptor Venus, resulting in a low FRET efficiency (Fig. 1 b). Upon activation of endogenous GEF by extracellular signals, GEF facilitates the release of GDP from Rac1, thereby converting Rac1 into a nucleotide-free form. Owing to the high intracellular concentration of GTP, Rac1 is then converted to the active form after autonomously binding to GTP. Intramolecular binding of active GTP-Rac1 to PAK CRIB brings CFP closer to Venus, thus enabling FRET from CFP to Venus to occur. The resulting Venus fluorescence provides an estimate of the activation state of Rac1 *in vivo*, with low and high ratios of Venus/CFP fluorescence corresponding to low and high levels of Rac1 activation, respectively.

We have previously revealed that the small GTPase OsRac1 is an important regulator controlling rice immunity (*15*, *16*, *26*), and monitoring its activation within living cells is therefore the next key step in further elucidating how plants trigger immunity. To monitor activation states of OsRac1, we have developed a plant version of the Raichu-Rac1system by combining the modification of human Raichu-Rac1 and a rice protoplast transfection system. Protoplasts do not possess a cell wall, and this enables direct live imaging of events both within the cell and at the cell surface, simultaneously and with no time delay in the response. Rice protoplasts also display a high growth rate and a high transfection rate, and we can control the expression levels of FRET sensors in plant cells without difficulty. Our work has pioneered the monitoring of spatiotemporal activities of plant small GTPases in living cells, which had been impossible by conventional biochemical methods (*19*). We have observed the resistance (R) protein Pit, an immune receptor that activates OsRac1 at the plasma membrane (*19*) (*27*), and reported that OsRac1 is activated within 3 min after treatment with a fungal cell wall component, chitin (*28*), which is a microbe-associated molecular pattern (MAMP) that triggers plant immunity. Moreover, by replacing the PAK CRIB domain in Raichu-OsRac1 with the N-terminal region of OsRbohB, an effector protein of OsRac1, we monitored the interaction between OsRac1 and OsRbohB and proposed that cytosolic Ca^2+^ concentration regulates the Rac-Rboh interaction in a dynamic manner (*29*). Since the principle of the intramolecular FRET biosensor appears to have universal applications for all of the small GTPases, we believe this Raichu system will become widely used in diverse research areas for understanding the spatiotemporal regulation of plant small GTPases ant this protocol can be widely adapted for for other FRET sensors in various plants.

## MATERIALS

- 2,4-dichlorophenoxyacetic acid (2,4-D) (C_8_H_6_Cl_2_O) (Sigma, cat. no. D7299)
- Agarose, low gelling temperature (Sigma, cat. no. A9414)
- Ammonium sulphate ((NH_4_)_2_SO_4_) (Sigma, cat. no. A3920)
- Boric acid (H_3_BO_3_) (Sigma, cat. no. B6768)
- Calcium chloride dihydrate (CaCl_2_•2H_2_O) (Sigma, cat. no. C5080)
- Cellulase RS (Yakult, cat. no. C8260)
- Chitin (N,N',N'',N''',N'''',N'''''-hexaacetylchitohexaose) (Carbosynth., cat. no. OH07433)
- Copper (II) sulphate pentahydrate (CuSO_4_•5H_2_O) (Sigma, cat. no. C8027)
- D-mannitol (C_6_H_14_O) (Sigma, cat. no. M1902)
- Deionized, distilled water (ddH_2_O)
- Ethylenediaminetetraacetic acid disodium salt dihydrate (Na_2_EDTA) (Sigma, cat. no. E5134)
- Glycine (C_2_H_5_NO) (Sigma, cat. no. G7126)
- Iron (II) sulphate heptahydrate (FeSO_4_•7H_2_O) (Sigma, cat. no. F8633)
- Macerozyme R10 (Yakult, cat. no. C1003)
- Magnesium chloride (MgCl_2_) (Sigma, cat. no. M8266)
- Magnesium sulphate heptahydrate (MgSO_4_•7H_2_O) (Sigma, cat. no. 63138)
- Manganese (II) sulphate tetrahydrate (MnSO_4_•4H_2_O) (Alfa Aesar, cat. no. B22081) ! CAUTION It is toxic and dangerous for the environment. Handle it with care, and wear gloves and eye protection.
- MES monohydrate (C_6_H_13_NO_4_S•H_2_O) (Fluka, cat. no. 69889)
- Murashige and Skoog (MS) Vitamin (Sigma, cat. no. M3900)
- Myo-inositol (C_6_H_12_O_6_) (Sigma, cat. no. I7508)
- Nicotinic acid (C_6_H_5_NO_2_) (Sigma, cat. no. N0761)
- Polyethylene glycol (PEG 4000) (Sigma, cat. no. 81240)
- Potassium chloride (KCl) (Sigma, cat. no. P9541)
- Potassium nitrate (KNO_3_) (Sigma, cat. no. P6083)
- Pyridoxine hydrochloride (C_8_H_12_ClNO_3_) (Sigma, cat. no. 47862)
- Plasmid Maxi Kit (Qiagen, cat. no. 12165)
- Sodium chloride (NaCl) (Sigma, cat. no. S5886)
- Sodium hydroxide (NaOH) (Sigma, cat. no. S8045) ! CAUTION NaOH is corrosive. Handle it with care, and wear gloves and eye protection.
- Sodium molybdate dihydrate (Na_2_MoO_4_•2H_2_O) (Sigma, cat. no. M1003)
- Sodium phosphate monobasic dihydrate (NaH_2_PO_4_•2H_2_O) (Sigma, cat. no. 71505)
- Sucrose (C_12_H_22_O_11_) (Sigma, cat. no. S0389)
- Thiamine hydrochloride (C_12_H_18_Cl_2_N_4_OS) (Sigma, cat. no. T1270)
- Zinc sulphate heptahydrate (ZnSO_4_•7H_2_O) (Sigma, cat. no. Z0251)

## EQUIPMENTS

- 0.2-μm filter 250 ml (CORNING 431096)
- 440-nm diode laser (iFLEX 2000, Point Source)
- 60× oil-immersion objective lens (UPlanSApo 60×/1.42, Olympus)
- Centrifuge with a swinging bucket rotor (Eppendorf: 5804R and S-4-72)
- Confocal scanner (Yokogawa CSU22)
- Cooled charge-coupled device camera (Hamamatsu Photonics, EM-CCD C9100-02).
- Cover glass (Matsunami Glass, Small: 24 × 32 mm, 0.13-0.17 mm, C024321, Large: 24 × 60 mm, 0.13-0.17 mm, C024601)
- Dual-View image splitter (DualView: Optical Insights)
- FRET filter for dual-emission imaging (Omega Optical, CFP: 480 ± 15 nm and Venus: 535 ± 20 nm)
- Inverted fluorescence microscope (Olympus IX-81)
- MetaMorph software (Universal Imaging)
- Microscope slide printed with highly water-repellent mark (Matsunami Glass, cat. no. TF0215)
- Nylon mesh (40 μm, Kyoshin Rikoh, cat. no. PP-40n)

## RECIPES

**2,4-D, 20 mg/ml** Dissolve 0.2 g of 2,4-D in 10 ml of 70% (v/v) Ethanol. Store the solution at 4 °C in dark for up to 6 months.

**CaCl_2,_ 1M** Dissolve 14.7 g of CaCl_2_•2H_2_O in 100 ml of ddH_2_O and autoclave. Store it at RT for up to 12 months.

**Cellulase solution** Add 140 mg of CaCl_2_•2H_2_O, 0.3 g of MES•H_2_O, 12.0 g of cellulase RS and 3.0 g of Macerozyme R10 to 200 ml of ddH_2_O and stir gently for 10-30 min. Next, add 18.2 g of D-mannitol and adjust the final volume to 300 ml with ddH_2_O. Filter the solution through three layers of filter paper. Adjust the pH to 5.6 and sterilize the solution with a 0.2-μm filter. Finally, divide into 20-ml aliquots in 50-ml tubes and store at -20 °C.

**FeEDTA, 1M** Dissolve 375 mg of Na_2_EDTA and 275 mg of FeSO_4_•7H_2_O in 80 ml of ddH_2_O. Adjust the final volume to 100 ml. Store at 4 °C.

**KNO_3_, 40 mM** Dissolve 0.4 g of KNO_3_ in 100 ml of ddH_2_O. Store at RT for up to 6 months.

**Mannitol, 0.8 M** Dissolve 14.6 g of D-mannitol in 100 ml of ddH_2_O and autoclave. Store at RT for up to 6 months.

**MgSO_4,_ 1 mM** Dissolve 25 mg of MgSO_4_•7H_2_O in 100 ml of ddH_2_O and autoclave. Store at RT for up to 6 months.

**MMG solution** Dissolve 14.6 g of D-mannitol, 0.6 g of MgCl_2_ and 170 mg of MES •H_2_O in 160 ml of ddH_2_O. Adjust the pH to 5.7 with 5N NaOH and dilute to 200 ml with ddH_2_O. Autoclave and divide into 15-ml Falcon tubes. Store at RT for up to 1 month.

**NaH_2_PO_4_, 2 mM** Dissolve 31 mg of NaH_2_PO_4_•2H_2_O in 100 ml of ddH_2_O. Store at RT for up to 6 months.

**(NH_4_)_2_SO_4_**, **2.5 mM** Dissolve 33 mg of MgSO_4_•7H_2_O in 100 ml of ddH_2_O. Store at RT for up to 6 months.

**PEG solution** Prepare just before use. Dissolve 4.0 g of PEG4000 in 3 ml of ddH_2_O. Add 2.5 ml of 0.8 M D-mannitol and 1 ml of 1 M CaCl_2._ Adjust volume to 10 ml with ddH_2_O. Dissolve and sterilize with a 0.2-μm filter.

**R2 Macro 20x solution** Dissolve 80.0 g of KNO_3_, 6.7 g of (NH_4_)_2_SO_4_, 5.0 g of MgSO_4_•7H_2_O, 3.0 g of CaCl_2_•2H_2_O and 5.5 g of NaH_2_PO_4_•2H_2_O in 800 ml of ddH_2_O. Adjust the volume to 1 l with ddH_2_O. Store at 4 °C.

**R2 Micro 1000x solution** Dissolve 160 mg of MnSO_4_•4H_2_O, 220 mg of ZnSO_4_•7H_2_O, 13 mg of CuSO_4_•5H_2_O, 0.6 g of H_3_BO_3_ and 13 mg of Na_2_MoO_4_•2H_2_O in 800 ml of ddH_2_O. Adjust the volume to 1 l with ddH_2_O and store at 4 °C.

**R2S** Dissolve 30.0 g of sucrose in 800 ml of ddH_2_O. Add 50 ml of 20x R2 Macro, 2 ml of FeEDTA, 1 ml of 1000x R2 Micro, 1 ml of MS Vitamin and 0.1 ml of 2,4-D. Adjust the pH of the solution to 5.7 with NaOH, and the volume to 1 l with ddH_2_O. Autoclave and store this medium at 4 °C for up to 1 month.

**W5 solution** Dissolve 1.8 g of NaCl, 2.8 g of CaCl_2_•2H_2_O, 70 mg of KCl and 90 mg of MES•H_2_O in 160 ml of ddH_2_O. Adjust the pH of the solution to 5.7 with 5N NaOH, and the volume to 200 ml with ddH_2_O. Autoclave and store at RT for up to 1 month.

## INSTRUCTION

This protocol describes a practical method for monitoring the activity of small GTPases in rice protoplasts using the intramolecular FRET biosensor. One of the most important factors for obtaining successful results is to develop a wide dynamic range of FRET biosensors. Since there are no standard protocols for making reliable Raichu constructs, we refrain from describing a detailed protocol on the development of FRET sensors in this protocol. Establishing reliable Raichu constructs requires a process of trial and error with many factors such as the choice of fluorescent proteins, the length of linkers and the combination of small GTPase and small GTPase-binding domain. For more information about sensor development, we refer readers to several excellent reviews of animal systems (*23*, *25*, *30*, *31*). The protocol that follows consists of four steps: 1) cell culture and preparation of Oc cells; 2) isolation of protoplasts; 3) transfection of protoplasts with *Raichu-OsRac1* plasmids; and 4) FRET imaging and data processing (Fig. 2).

**Figure 2.**
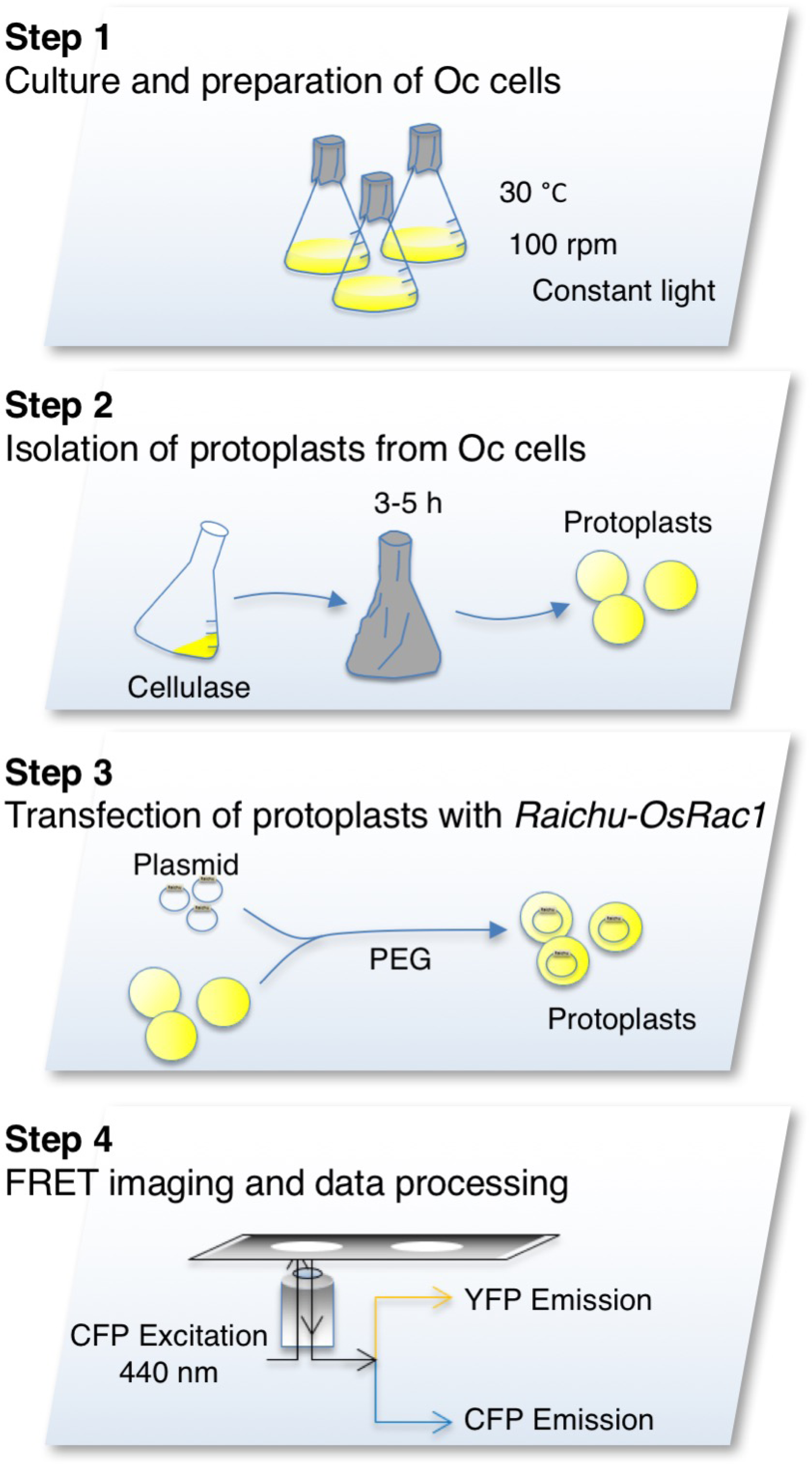
Experimental flow

### Construction of Raichu-OsRac1

We first tested the binding specificity of the CRIB domain of PAK for OsRac1 using a pull-down assay. PAK CRIB specifically bound to the constitutively active (G19V) form of OsRac1 (OsRac1 CA) and not to the dominant negative (T24N) form (OsRac1 DN), indicating that the interaction between PAK CRIB and OsRac1 occurs only when OsRac1 is activated (Fig. 1 b and c). The backbone of our Raichu-OsRac1 is essentially identical to that made by Itoh et al. (*24*), except that human Rac1 and the C-terminal polybasic region and CAAX box of human Ki-Ras are replaced by rice OsRac1 and the corresponding elements of OsRac1, respectively (*19*). The choice of C-terminal polybasic region and post-translational modification site is also important, because subcellular localization of small GTPases is strongly influenced by these functions (*29*). To determine whether the activation of OsRac1 increases FRET efficiency *in vivo*, we also prepared CA and DN forms of Raichu-OsRac1.

#### [1] Cell culture and preparation of Oc cells - Time: 3 days

We used *Oryza sativa* L. C5924 (Oc) cells to generate rice protoplasts for transfection assays of Raichu-OsRac1, because their transfection efficiency is much higher than that of protoplasts derived from other rice cultivars (*32*).

*All procedures should be done on a clean bench to avoid contamination

From a fully saturated 20-ml suspension of Oc cells, we discard all medium as well as about half of the cells, and then add 20 ml of fresh R2S medium in a 100-ml flask every week to maintain the cells. Cells are cultured at 30°C with rotation at 100 rpm under constant light. We change the flasks every month because debris accumulates on their inner surface.

1. **Use fully saturated suspension cells cultured in fresh R2S medium for 3 days.**

For transfection experiments, the medium of a fully saturated cell suspension is replaced with 20 ml of fresh R2S medium and the cells are cultured for 3 days.

#### [2] Isolation of protoplasts from Oc cells - Time: 6 h

We developed a rice protoplast isolation and transfection method, according to Yoo et al., with minor modifications (*33*). Cell walls of Oc cells are removed using cellulase and the protoplasts transfected with *Raichu-OsRac1* vectors by a PEG method.

*All procedures should be done on a clean bench to avoid contamination (Fig. 3 b).

**Figure 3.**
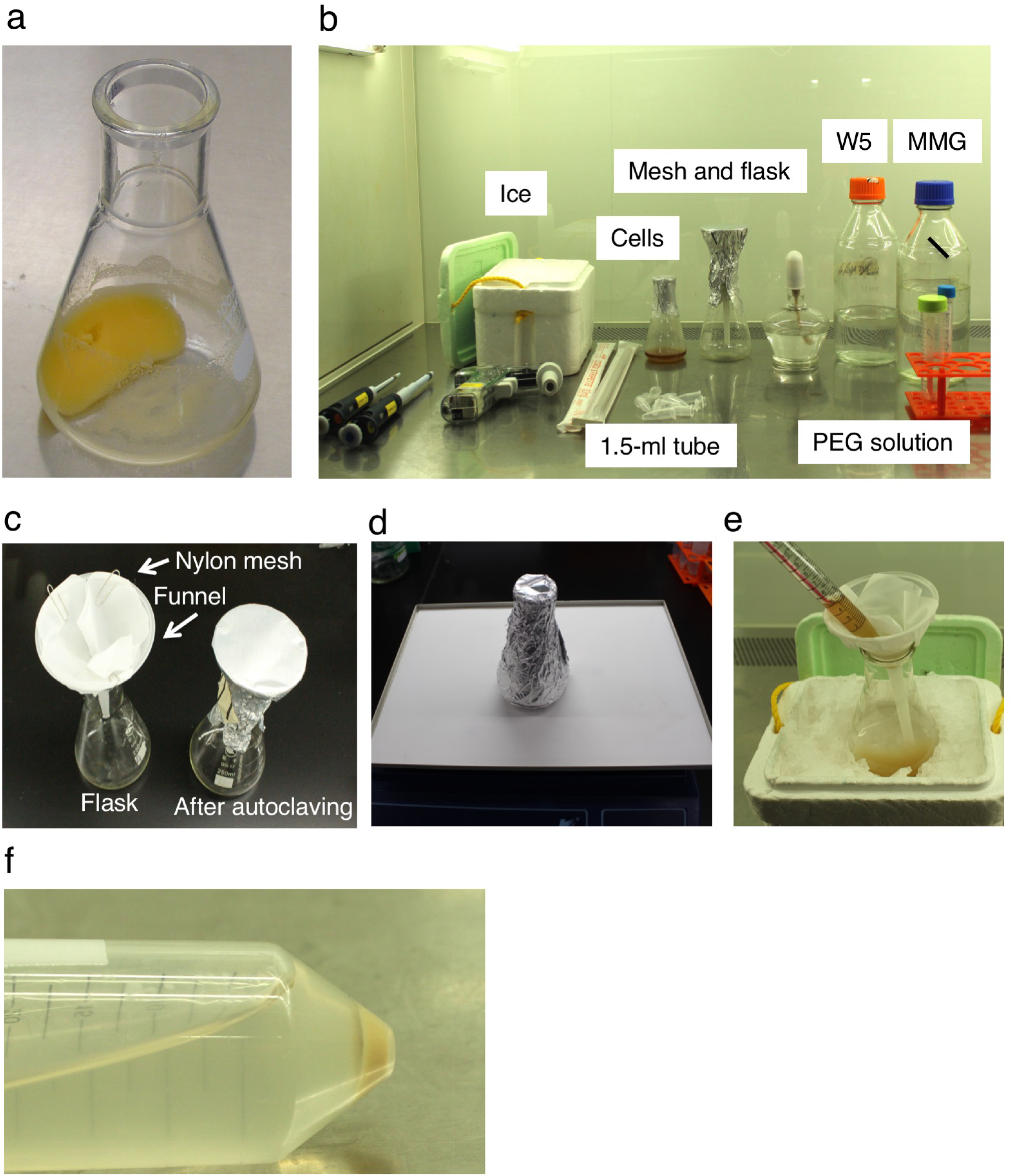
Materials for transfection of rice protoplasts with *Raichu-OsRac1*. **a,** Healthy Oc cells. **b,** Materials required for transfection on a clean bench. **c,** Preparation of a funnel with a nylon mesh filter. **d,** Cellulase treatment on an orbital shaker. Incubate the cells with shaking at 50 rpm. **e**, Filtration of cellulase-treated Oc cells. **f,** Precipitated cells after centrifugation with a swinging bucket rotor.

1. **Prepare a conical flask with a funnel and nylon mesh filter, and autoclave** (Fig. 3 c)
2. **Irradiate a clean bench with ultraviolet light for 10-20 min**
3. **Pre-warm cellulase solution at 30 °C to thaw**
4. **Completely remove R2S medium from a flask containing a 3-day culture** (Fig. 3 a); **add 20 ml of cellulase solution and cover the flask with aluminium foil**
5. **Incubate the cells at 30 °C with shaking at 50 rpm for 3-5 h** (Fig. 3 d)
6. **Add 20 ml of W5 solution and filter the cells through a sheet of 40-µm nylon mesh** (Fig. 3 e) * All procedures should be conducted on ice. To prevent the filters from clogging, we pipette the cells gently in 5-ml aliquots into the filter.
7. **Transfer the filtered cells to a 50-ml Falcon tube**
8. **Centrifuge the tube at 120 *g* for 4 min using a swinging bucket rotor** Check whether sufficient cells are pelleted in the bottom of the tube (Fig. 3 f).
9. **Decant and discard the supernatant**
10. **Add 20 ml of W5 solution to the protoplast pellet and suspend it by gentle swirling**
11. **Centrifuge at 120 *g* for 4 min using a swinging bucket rotor**
12. **Discard the supernatant and add 2 ml of W5 solution** To avoid cell damage, resuspend the cells gently.
13. **Keep the suspension on ice for 30 min**
14. **Dilute 10 μl of the protoplast suspension with 90 μl of W5 solution to measure the density of protoplasts using a haemocytometer, and estimate the total number of protoplasts** We count only intact, round cells. In general, we can obtain 2-8×10^6^ cells from one flask. If your yield is very low, start again from the beginning.
15. **Centrifuge at 120 *g* for 4 min using a swinging bucket rotor and discard the supernatant**
16. **Resuspend the protoplasts at 2×10^6^ cells/ml with MMG**

* CRITICAL STEP For general experiments such as localization studies and analyses of gene expression, protoplasts can be kept at 4 °C for up to 24 h, but freshly prepared protoplasts are essential for Raichu-OsRac1 experiments.

#### [3] Transfection of protoplasts with *Raichu-OsRac1* plasmids

In general, we are able to observe a 30-40% transfection efficiency with control green fluorescent protein (GFP), and it is very important to keep this efficiency high to obtain reproducible and reliable results. Since it is easy to generate CA and DN mutants of small GTPases, we strongly recommend adding *Raichu-OsRac1 CA* and *DN* vectors as positive and negative controls to ensure a wide dynamic range of the Venus/CFP ratio in each experiment.

* All steps are carried out at 25°C (room temperature).

* Prepare plasmid DNA for transfection and fresh PEG solution.

The concentration of all of plasmids is adjusted to 2 μg/μl to minimize the volume of plasmid used in the transfection.

1. **Add the required plasmids to a 1.5-ml tube** * CRITICAL STEP It is essential to optimize the amount and ratio of plasmids to suit your experimental design. The total amount of plasmid in each sample should not exceed 10 μl or 10 μg, and should be the same for both test and control vectors.
2. **Add 100 µl of protoplast suspension (2×10^5^ cells)**
3. **Add 110 µl of PEG solution and mix gently by inverting the tube five times**
4. **Incubate the transfection mixture for 20 min at room temperature**
5. **Add 1 ml of W5 solution to the tube and mix gently by inverting five times**
6. **Centrifuge at 120 *g* for 10 min and discard the supernatant**
7. **Add 100 µl of W5 solution**
8. **Suspend the protoplasts gently and place the tube in a light-resistant box on a slant** We put the tube on a slant to prevent the fragile protoplasts from settling in the bottom of the tube, and we use a light-resistant box to prevent bleaching of the fluorescent proteins.
9. **Incubate in the dark at 30 °C for 10-16 h**

#### [4] FRET imaging and data processing

Raichu-OsRac1 proteins begin to be expressed 8 h after transfection, and it usually takes until about 10 h after transfection to achieve sufficient expression of the biosensors. The optimal observation time is 10-16 h after transfection. FRET efficiencies may vary depending on the timing of observation, due to differences in the time required for maturation of donor and acceptor fluorescent proteins, and it is better to avoid observing cells beyond 16 h after transfection in our condition. The expression levels of Raichu-OsRac1 are different in individual cells and it is important to select cells displaying appropriate localization (*34*) and comparable levels of Raichu-OsRac1 (*19*).

To obtain images of Raichu-OsRac1, we use an inverted microscope equipped with a Confocal Scanner Unit, a Dual-View image splitter and a CCD camera, which can simultaneously take CFP and Venus images. The donor protein CFP is excited by a 440-nm diode laser. Average values for the fluorescence intensity of CFP and Venus in the region of interest are calculated after subtracting background fluorescence. The normalized FRET efficiency is then calculated according to Sorkin et al. (*35*).

1. **Mount 10 μl of the transfected rice protoplasts on a microscope slide** We keep the 1.5-ml tube upright for 30 min to collect the transfected cells at the bottom of the tube, and then transfer 10 μl of the cells to the recess of a microscope slide glass printed with a highly water-repellent mark and cover it with a cover glass using a nail varnish (Fig. 4 a).

**Figure 4.**
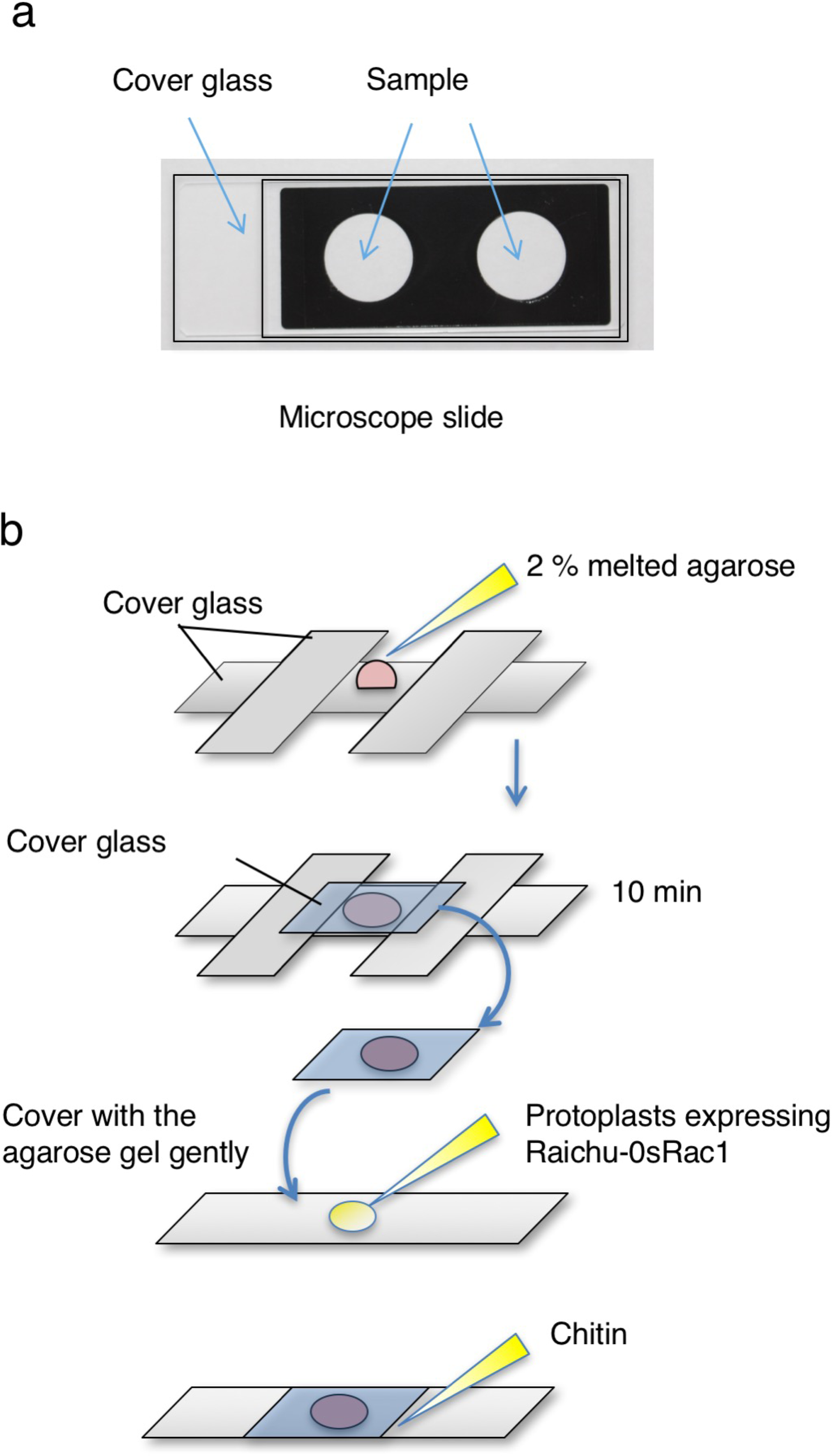
Mounting the transfected rice protoplasts on a microscope slide. **a,** Mounting transfected cells on a microscope slide. **b,** Preparing a cover glass coated In time-lapse experiments using chitin, we prepare a smaller cover glass (24 × 32 mm, thickness 0.12-0.17 mm) coated on its underside with 2% low-melting-temperature agarose, according to Fig. 4 b. We sandwich the melted agarose between the smaller and a larger (24 × 60 mm, thickness 0.12-0.17 mm) cover glass, using two large cover glasses as spacers, and allow it to set for 10 min. We put 10 μl of the transfected cells onto a new larger cover glass. To situate the cells, they are sandwiched between this larger cover glass and the smaller cover glass coated with low-melting-temperature agarose. Chitin is added via a small opening between the two cover glasses.
2. **Open MetaMorph software and obtain images of Raichu-OsRac1** We use an inverted microscope (IX81: Olympus) equipped with a Confocal Scanner Unit (CSU22: Yokokawa), a Dual-View image splitter (DualView: Optical Insights), and a CCD camera (EM-CCD C9100-02: Hamamatsu). MetaMorph is used for capturing and analysing images. Using Multi Dimensional Acquisition Mode, donor protein CFP is excited by a 440-nm diode laser, and we then take images for CFP (480 ± 15 nm) and Venus (535 ± 20 nm) at the same time. Our typical imaging conditions are sensitivity 50-150, and exposure time 1000-1500 msec. It is important to obtain images from healthy transfected cells that show appropriate localization of Raichu-OsRac1 in the plasma membrane (*34*), and also to carefully adjust sensitivity and exposure time to avoid saturation of the fluorescence images. Finally, we take photos of the protoplasts expressing Raichu-OsRac1.
3. **Subtract background and make ratio images** The emission spectra of CFP and Venus fluorescence are obtained simultaneously using a Dual-View image splitter. This slightly misaligns the position of CFP and Venus images, and the misalignment is corrected using the Align function of MetaMorph software. To precisely measure the fluorescence intensities of CFP and Venus, background fluorescence is subtracted from the images. The Region of Interest (ROI) is selected within appropriate areas that are devoid of fluorescent objects, and this background fluorescent intensity is subtracted from whole images.
4. **Merge processed CFP and Venus images into a single image and measure the Venus/CFP ratio** Using the background-subtracted CFP and Venus images, Venus images are divided by CFP images, thereby creating Venus/CFP ratio images. To quantitatively measure the Venus/CFP ratio images, we set ROIs over the cells of interest to measure the average intensity of Venus and CFP images using the Region Measurement function of MetaMorph software, and the obtained value is exported to a Microsoft Excel file. To visualize the activation level of OsRac1, the Venus/CFP ratio images are pesudo-coloured with the intensity-modulated display (IMD) mode, which divides the mean signal intensity of the Venus/CFP ratio into eight colours from red to blue (Fig. 5). In general, we can obtain sufficient dynamic ranges when we set the upper and lower limits of the IMD mode at about 0.5 higher and lower than the average ratio of Venus/CFP for Raichu-CA-OsRac1 and -DN-OsRac1, respectively.

**Figure 5.**
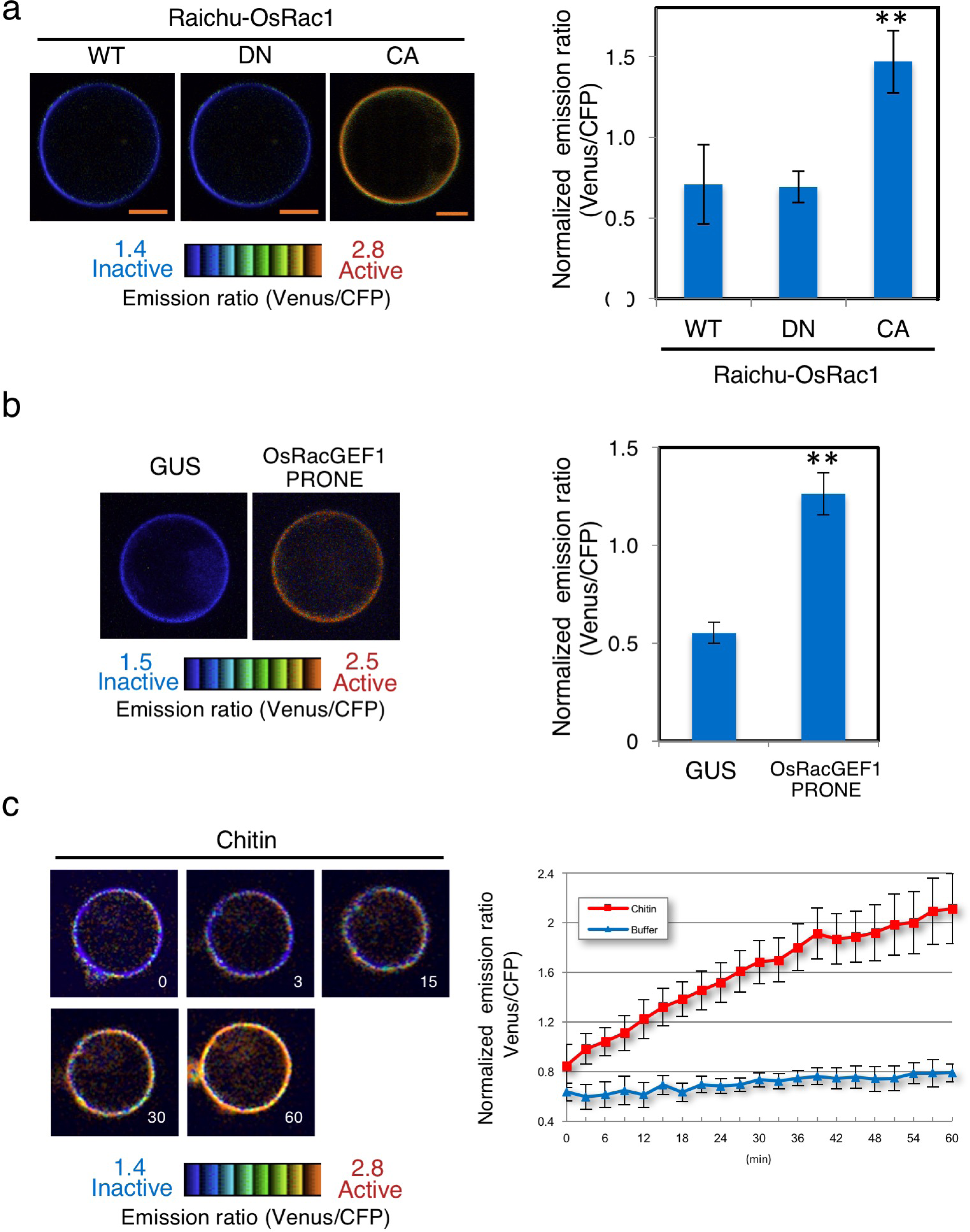
Validating the Raichu-OsRac1 FRET sensor and monitoring OsRac1 activation after treatment of rice protoplasts with the MAMP chitin. **a and b,** Emission ratio images of rice protoplasts expressing Raichu-OsRac1 mutants. Rice protoplasts were transfected with constructs expressing Raichu-OsRac1 WT, mutants and the PRONE domain of OsRacGEF1. Double asterisks indicate significant differences from the data for WT-OsRac1 or control GUS enzyme (*P* < 0.01). Error bars indicate SE (n > 30). c, Time-lapse imaging of rice protoplasts expressing Raichu-OsRac1-WT after chitin treatment. Red line, 0.5 µg/ml chitin treatment; blue line, W5 buffer treatment. The FRET images are shown in IMD mode, which associates colour hue with emission ratio values and the intensity of each hue with the brightness of the source image. Bars: 5 µm.
5. **Process data** For statistical analyses, **corrected FRET (cFRET)** followed by **normalized FRET (nFRET)** are calculated, according to Sorkin et al., (*35*) with minor modifications. Fluorescence in the Venus channel consists of FRET and non-FRET fluorescence derived from the cross-over of Venus and CFP fluorescence through the FRET filters. Cross-over between Venus and CFP fluorescence divides into two factors: CFP emission detected in the Venus channel (xCFP) and Venus emission induced by CFP excitation (yVenus). cFRET is calculated as follows: **cFRET = total Venus – xCFP – yVenus**. The extent of cross-over (x and y) is a characteristic value in each optical system and must be obtained in advance using cells expressing either Venus or CFP and the following formulae: Cells expressing CFP x = Fluorescence in Venus channel when CFP is activated by CFP excitation / CFP emission induced by CFP excitation Cells expressing Venus y = Fluorescence in Venus channel when Venus is activated by CFP excitation / Venus emission induced by Venus excitation In our optical system, x and y are about 0.77 and 0.02, respectively, which are conservative values that we can continue to use until we change the optical system. The y value is much smaller than the x value, and we do not take the y value into consideration. As shown in the formula above, the cFRET value is affected by the expression level of CFP and Venus; thus, cFRET is divided by CFP as follows: **nFRET = cFRET/CFP emission induced by CFP excitation.** To examine whether the activation of OsRac1 increased FRET efficiency in living cells, we transfected rice protoplasts with Raichu-OsRac1 WT, DN and CA. OsRac1 has been shown previously to localize to the plasma membrane (*34*). Consistent with this, Raichu-OsRac1 was localized mainly at the plasma membrane. The ratio of Venus/CFP fluorescence in protoplasts expressing Raichu-CA-OsRac1 was higher than those in protoplasts expressing either Raichu-WT-OsRac1 or Raichu-DN-OsRac1 (Fig. 5 a), indicating that the interaction between CRIB and OsRac1 occurs only when OsRac1 is activated. These results demonstrate that the ratios of Venus/CFP fluorescence for Raichu-OsRac1 reflect the activation state of OsRac1 in rice protoplasts.

We have previously found that a PRONE-type GEF, OsRacGEF1, displays GEF activity towards OsRac1 *in vitro* and plays an important role in the activation of OsRac1 in chitin-induced immune responses (*28*). To extend that observation, we next tried to monitor the activation of OsRac1 by OsRacGEF1 in living cells. In control GUS-transfected cells, the ratio of Venus/CFP fluorescence of Raichu-OsRac1 was low, but when we expressed the PRONE domain of OsRac1GEF1 (OsRacGEF1 PRONE), the ratio increased (Fig. 5 b), showing that OsRacGEF1 PRONE indeed activates OsRac1 in rice protoplasts.

To elucidate the spatiotemporal regulation of OsRac1 activation in MAMP-triggered immunity, we treated the rice protoplasts with chitin, which is a component of pathogenic and non-pathogenic fungi and is one of the best-studied MAMPs. Chitin markedly and rapidly induced OsRac1 activation in the rice protoplasts; in contrast, the OsRac1 activation level was negligible in cells treated with buffer (Fig. 5 c). Together, these results show that OsRac1 is rapidly activated at the plasma membrane of rice protoplasts after chitin elicitor treatment.

### **TROUBLESSHOOTING** (Also see Table 1)

The condition of the Oc cells is the most important factor to obtain successful results, and the cells have to be treated carefully. Cells that are in good condition are bright yellow, and a white colour indicates that they are not in good condition (Fig. 3 a). The incubation time strongly affects transfection efficiency and yield of cells. In Oc cells, a 3-h treatment is sufficient to obtain protoplasts, but if you cannot obtain sufficient protoplasts, you can extend the incubation time to 5 h. If you are using different rice cultivars, it may be desirable to optimize the incubation time for your experimental conditions in advance.

**Table 1.** Troubleshooting

| Problem | Possible reason | Solution |
| --- | --- | --- |
| Low protoplast yield and low transfection efficiency | Condition of suspension cells | Suspension cell condition is the most important factor for obtaining successful results. If you have problems with a transfection, prepare new suspension cells from your stock or rice seeds |
|  | Insufficient cellulase digestion | Optimize the period of cellulase treatment or prepare fresh cellulase solution |
|  | Problem with transfection reagents | Prepare new transfection reagents. PEG solution can not be stored, and you should prepare fresh PEG for each experiment |
|  | Low DNA quality | Use an animal transfection grade of DNA purification kit |
| Decay of FRET signal during imaging | Photobleaching by the excitation | Minimize exposure time and duration of laser excitation |

Plasmid purity affects the transfection efficiency and we highly recommend the use of animal transfection grade plasmid purification kits or CsCl gradient purification.

## NOTES and REMARKS

It was important for us to address the issue of whether OsRac1 is activated following pathogen recognition *in planta*. We therefore tried to make transgenic rice plants expressing Raichu-OsRac1. Unfortunately, we have not yet been able to observe Raichu-OsRac1 fluorescence *in planta*. This may be due to homology-dependent silencing of YFP and CFP expression, a problem that has been encountered by other researchers endeavouring to develop stably expressed FRET biosensors *in planta* (*36*). Further work is necessary to resolve this issue.

## ACKNOELEDGEMENTS

We thank Dr. Michiyuki Matsuda for providing the Raichu-Rac1 vector. We appreciate Ms. Tomoko Aoi and Ms. Xiaoyun Fang for excellent technical assistance. We also thank the members of the Laboratory of Plant Molecular Genetics at NAIST and the Laboratory of Signal Transduction and Immunity at PSC for invaluable support and discussions. A.A. was supported by a fellowship from JSPS. Y.K. was supported by JSPS KAKENHI, the Takeda Science Foundation, Chinese Academy of Sciences, Shanghai Institutes for Biological Sciences, Shanghai Center for Plant Stress Biology, National Natural Science Foundation in China, and Chinese Academy of Sciences Hundred Talents Program.

H. L. W., A. A, Y. K., and K. S. designed the study; H. L. W., A. A., M. H., T. M., J. O., K. K., Q. W. and Y. K. performed experiments and analyzed data; Y. K. wrote the manuscript; M. H., N. I., T. K., and K. S. gave technical support and conceptual advice.

## REFERENCES

1. R. Y. Tsien, A. Miyawaki, Seeing the machinery of live cells. Science 280, 1954–1955 (1998).

2. E. A. Jares-Erijman, T. M. Jovin, FRET imaging. Nat Biotechnol 21, 1387–1395 (2003).

3. K. Aoki, M. Matsuda, Visualization of small GTPase activity with fluorescence resonance energy transfer-based biosensors. Nature protocols 4, 1623–1631 (2009).

4. A. Miyawaki, J. Llopis, R. Heim, J. M. McCaffery, J. A. Adams, M. Ikura, R. Y. Tsien, Fluorescent indicators for Ca2+ based on green fluorescent proteins and calmodulin. Nature 388, 882–887 (1997).

5. N. Mochizuki, S. Yamashita, K. Kurokawa, Y. Ohba, T. Nagai, A. Miyawaki, M. Matsuda, Spatio-temporal images of growth-factor-induced activation of Ras and Rap1. Nature 411, 1065–1068 (2001).

6. F. Sipieter, P. Vandame, C. Spriet, A. Leray, P. Vincent, D. Trinel, J. F. Bodart, F. B. Riquet, L. Heliot, From FRET imaging to practical methodology for kinase activity sensing in living cells. Prog Mol Biol Transl Sci 113, 145–216 (2013).

7. M. Paduch, F. Jelen, J. Otlewski, Structure of small G proteins and their regulators. Acta Biochim Pol 48, 829–850 (2001).

8. K. Wennerberg, K. L. Rossman, C. J. Der, The Ras superfamily at a glance. J Cell Sci 118, 843–846 (2005).

9. H. Li, G. Wu, D. Ware, K. R. Davis, Z. Yang, Arabidopsis Rho-related GTPases: differential gene expression in pollen and polar localization in fission yeast. Plant Physiol 118, 407–417 (1998).

10. T. Kawasaki, K. Henmi, E. Ono, S. Hatakeyama, M. Iwano, H. Satoh, K. Shimamoto, The small GTP-binding protein Rac is a regulator of cell death in plants. Proc Natl Acad Sci U S A 96, 10922–10926 (1999).

11. P. Winge, T. Brembu, R. Kristensen, A. M. Bones, Genetic structure and evolution of RAC-GTPases in Arabidopsis thaliana. Genetics 156, 1959–1971 (2000).

12. J. Cherfils, M. Zeghouf, Regulation of small GTPases by GEFs, GAPs, and GDIs. Physiol Rev 93, 269–309 (2013).

13. C. Craddock, I. Lavagi, Z. Yang, New insights into Rho signaling from plant ROP/Rac GTPases. Trends Cell Biol 22, 492–501 (2012).

14. S. Yalovsky, Protein lipid modifications and the regulation of ROP GTPase function. J Exp Bot 66, 1617–1624 (2015).

15. Y. Kawano, T. Kaneko-Kawano, K. Shimamoto, Rho family GTPase-dependent immunity in plants and animals. Frontiers in plant science 5, 522 (2014).

16. Y. Kawano, K. Shimamoto, Early signaling network in rice PRR- and R-mediated immunity. Curr Opin Plant Biol. 16, 496–504 (2013).

17. E. E. Sander, S. van Delft, J. P. ten Klooster, T. Reid, R. A. van der Kammen, F. Michiels, J. G. Collard, Matrix-dependent Tiam1/Rac signaling in epithelial cells promotes either cell-cell adhesion or cell migration and is regulated by phosphatidylinositol 3-kinase. J Cell Biol 143, 1385–1398 (1998).

18. L. Z. Tao, A. Y. Cheung, H. M. Wu, Plant Rac-like GTPases are activated by auxin and mediate auxin-responsive gene expression. Plant Cell 14, 2745–2760 (2002).

19. Y. Kawano, A. Akamatsu, K. Hayashi, Y. Housen, J. Okuda, A. Yao, A. Nakashima, H. Takahashi, H. Yoshida, H. L. Wong, T. Kawasaki, K. Shimamoto, Activation of a Rac GTPase by the NLR family disease resistance protein Pit plays a critical role in rice innate immunity. Cell Host Microbe 7, 362–375 (2010).

20. T. Xu, M. Wen, S. Nagawa, Y. Fu, J. G. Chen, M. J. Wu, C. Perrot-Rechenmann, J. Friml, A. M. Jones, Z. Yang, Cell surface- and rho GTPase-based auxin signaling controls cellular interdigitation in Arabidopsis. Cell 143, 99–110 (2010).

21. V. S. Kraynov, C. Chamberlain, G. M. Bokoch, M. A. Schwartz, S. Slabaugh, K. M. Hahn, Localized Rac activation dynamics visualized in living cells. Science 290, 333–337 (2000).

22. Y. Fu, Y. Gu, Z. Zheng, G. Wasteneys, Z. Yang, Arabidopsis interdigitating cell growth requires two antagonistic pathways with opposing action on cell morphogenesis. Cell 120, 687–700 (2005).

23. A. Miyawaki, Visualization of the spatial and temporal dynamics of intracellular signaling. Dev Cell 4, 295–305 (2003).

24. R. E. Itoh, K. Kurokawa, Y. Ohba, H. Yoshizaki, N. Mochizuki, M. Matsuda, Activation of rac and cdc42 video imaged by fluorescent resonance energy transfer-based single-molecule probes in the membrane of living cells. Mol Cell Biol 22, 6582–6591 (2002).

25. E. Kiyokawa, K. Aoki, T. Nakamura, M. Matsuda, Spatiotemporal regulation of small GTPases as revealed by probes based on the principle of Forster Resonance Energy Transfer (FRET): Implications for signaling and pharmacology. Annual review of pharmacology and toxicology 51, 337–358 (2011).

26. Y. Kawano, L. Chen, K. Shimamoto, The function of Rac small GTPase and associated proteins in rice innate immunity. Rice 3, 112–121 (2010).

27. Y. Kawano, T. Fujiwara, A. Yao, Y. Housen, K. Hayashi, K. Shimamoto, Palmitoylation-dependent membrane localization of the rice resistance protein pit is critical for the activation of the small GTPase OsRac1. J Biol Chem 289, 19079–19088 (2014).

28. A. Akamatsu, H. Wong, M. Fujiwara, J. Okuda, K. Nishide, K. Uno, K. Imai, K. Umemura, T. Kawasaki, Y. Kawano, K. Shimamoto, An OsCEBiP/OsCERK1-OsRacGEF1-OsRac1 module is an essential component of chitin-induced rice immunity. Cell Host Microbe 13, 465–476 (2013).

29. H. L. Wong, R. Pinontoan, K. Hayashi, R. Tabata, T. Yaeno, K. Hasegawa, C. Kojima, H. Yoshioka, K. Iba, T. Kawasaki, K. Shimamoto, Regulation of rice NADPH oxidase by binding of Rac GTPase to its N-terminal extension. Plant Cell 19, 4022–4034 (2007).

30. T. Nakamura, K. Aoki, M. Matsuda, Monitoring spatio-temporal regulation of Ras and Rho GTPase with GFP-based FRET probes. Methods 37, 146–153 (2005).

31. N. Komatsu, K. Aoki, M. Yamada, H. Yukinaga, Y. Fujita, Y. Kamioka, M. Matsuda, Development of an optimized backbone of FRET biosensors for kinases and GTPases. Mol Biol Cell 22, 4647–4656 (2011).

32. A. Baba, S. Hasezawa, Syono K, Cultivation of Rice Protoplasts and Their Transformation Mediated by Agrobacterium Spheroplasts. Plant Cell Physiol 27, 463–471 (1986).

33. S. D. Yoo, Y. H. Cho, J. Sheen, Arabidopsis mesophyll protoplasts: a versatile cell system for transient gene expression analysis. Nature protocols 2, 1565–1572 (2007).

34. E. Ono, H. L. Wong, T. Kawasaki, M. Hasegawa, O. Kodama, K. Shimamoto, Essential role of the small GTPase Rac in disease resistance of rice. Proc Natl Acad Sci U S A 98, 759–764 (2001).

35. A. Sorkin, M. McClure, F. Huang, R. Carter, Interaction of EGF receptor and grb2 in living cells visualized by fluorescence resonance energy transfer (FRET) microscopy. Curr Biol 10, 1395–1398 (2000).

36. K. Deuschle, B. Chaudhuri, S. Okumoto, I. Lager, S. Lalonde, W. B. Frommer, Rapid metabolism of glucose detected with FRET glucose nanosensors in epidermal cells and intact roots of Arabidopsis RNA-silencing mutants. Plant Cell 18, 2314–2325 (2006).

